# OTX2 homeoprotein functions in adult choroid plexus

**DOI:** 10.1101/2021.04.28.441734

**Authors:** Anabelle Planques, Vanessa Oliveira Moreira, David Benacom, Clémence Bernard, Laurent Jourdren, Corinne Blugeon, Florent Dingli, Vanessa Masson, Damarys Loew, Alain Prochiantz, Ariel A. Di Nardo

**Affiliations:** Centre for Interdisciplinary Research in Biology (CIRB), Collège de France, CNRS UMR7241, INSERM U1050, Labex MemoLife, PSL University, Paris, France; Genomics Core Facility, Institut de Biologie de l’ENS (IBENS), Département de Biologie, École Normale Supérieure, CNRS, INSERM, PSL University, 75005 Paris, France; Institut Curie, Centre de Recherche, Laboratoire de Spectrométrie de Masse Protéomique, 75248 Paris Cedex 05, France; Institute of Neurosciences, Chinese Academy of Sciences, 320 Yue Yang Road, Shanghai, 200031, China

**Keywords:** homeodomain, transcription factor, splicing, homeostasis

## Abstract

Choroid plexus secretes cerebrospinal fluid important for brain development and homeostasis. The OTX2 homeoprotein is critical for choroid plexus development and remains highly expressed in adult choroid plexus. Through RNA sequencing analyses of constitutive and conditional knockdown adult mouse models, we reveal putative roles for OTX2 in choroid plexus function, including cell signaling and adhesion, and show that it regulates the expression of factors secreted into cerebrospinal fluid, notably transthyretin. We show that *Otx2* expression impacts choroid plexus immune and stress responses, and also affects splicing which leads to changes in mRNA isoforms of proteins implicated in oxidative stress response and DNA repair. Through mass spectrometry analysis of OTX2 protein partners in the choroid plexus, and in known non-cell autonomous target regions such as visual cortex and ventricular-subventricular zone, we identified putative targets involved in cell adhesion, chromatin structure and RNA processing. Thus, OTX2 retains important roles in choroid plexus function and brain homeostasis throughout life.

## Introduction

Choroid plexus (ChP) epithelium is located in the brain ventricles and secretes cerebrospinal fluid (CSF) containing molecules that regulate embryonic brain development and adult brain homeostasis (Fame and Lehtinen, 2020). The ventricular system includes the two lateral ventricles (LVs) in each cerebellar hemisphere, the central third ventricle of the forebrain diencephalon, and the central fourth ventricle (4V) in the hindbrain. This system is interconnected allowing for CSF flow throughout and is also connected via the 4V with the central canal of the spinal cord. The OTX2 homeoprotein transcription factor is critical for ChP embryonic development and functions (Johansson et al., 2013). Interestingly, temporal and spatial heterogeneity is evident, as the role of OTX2 evolves during development and differs between ChPs. For example, in late embryonic development OTX2 is required for the maintenance of the 4V ChP but not LV ChP (Johansson et al., 2013). Indeed, embryonic LV and 4V ChP show distinct gene expression patterns (Lun et al., 2015), suggesting different signaling properties. In the adult, OTX2 is still strongly expressed by the ChP (Spatazza et al., 2013), but its role has not been thoroughly investigated (Spatazza et al., 2013).

Homeoproteins are transcription factors important for embryonic development and adult homeostasis and cell survival, and several homeoproteins have been shown to have functions beyond transcription including translation regulation, DNA repair and signal transduction (Di Nardo et al., 2018; Rezsohazy, 2014). While several studies have explored the molecular partners and transcriptional targets of OTX2, they were typically restricted to embryonic contexts (Beby and Lamonerie, 2013; Fossat et al., 2006; Hoch et al., 2015). In the adult mouse, recent analyses of OTX2 protein and DNA targets have been performed on retina (Fant et al., 2015; Samuel et al., 2014). These studies revealed targets implicated in tissue development and homeostasis, and confirmed that OTX2 not only binds multiple sites across DNA, but also interacts with the machinery for RNA processing, export, and translation. To examine the role of OTX2 in adult ChP, we use a mouse model for constitutive heterozygous *Otx2* knockdown and a model for ChP-specific conditional knockdown of *Otx2*. Through transcriptomic analysis of LV and 4V ChPs, we reveal dysregulation of cell adhesion and membrane proteins, secreted factors, signaling factors, immune response and oxidative stress response. OTX2 is not only highly expressed in the ChP, it is also secreted into CSF and distributed throughout various neurogenic niches and cortical structures, with an identified role in regulating plasticity of neural circuits implicated in sensory processing and cognition (Di Nardo et al., 2020). Through mass spectrometry analysis of OTX2 partners in ChP and in non-cell autonomous target regions (Planques et al., 2019; Sugiyama et al., 2008), including ventricular-subventricular zone (V-SVZ), rostral migratory stream (RMS) and visual cortex (VCx), we identified putative targets and interactors involved in cell adhesion, chromatin structure and RNA processing. We also performed splice variant analysis and confirmed by acute *Otx2* ChP knockdown in adult wildtype mice that OTX2 can regulate isoform distribution of genes involved in stress response and DNA repair. Taken together, our findings suggest that OTX2 has direct roles in ChP signaling, barrier, and surveillance functions.

## Results and Discussion

### Conditional and constitutive knockdown of Otx2 in adult ChP

OTX2 is a key regulator of ChP and brain development but its role in adult ChP is not well known. To gain insight into its adult “ChP functions”, we performed RNA sequencing analysis with two mouse models. The first consisted of 3-month-old *Otx2^lox/lox^* mice for conditional knockdown of *Otx2* specifically in the ChP through intracerebroventricular (icv) injections of Cre-Tat recombinant protein (Spatazza et al., 2013). The ChPs from LV and 4V were dissected separately from both Cre-Tat injected (Cre^+^*Otx2^lox/lox^*) and control vehicle-injected mice (Veh^+^*Otx2^lox/lox^*). While the bilateral stereotaxic injections of vehicle or Cre-Tat are performed only in lateral ventricles, we have previously shown that the level of *Otx2* knockdown in 4V ChP is proportional to that in LV ChP (Planques et al., 2019). The second model consisted of *Otx2^+/GFP^* mice as a constitutive heterozygous knockout mutant with ~50% Otx2 protein levels compared to wildtype (Bernard et al., 2014). For this model, only the 4V ChPs were dissected and pooled from 3-month-old wildtype and mutant mice.

The transcriptomics analysis of adult ChP showed highly expressed genes involved in energy metabolism, protein signaling, solute transport, cell adhesion, cytoskeleton and chaperone activity (Table 1). While not in exactly the same order of gene expression level, this list compares favorably with those obtained from other ChP transcriptomics studies (Baruch et al., 2014; Silva-Vargas et al., 2016). The conditional adult mouse knockdown of *Otx2* led to significant changes in expression of 375 genes in LV ChP and 808 genes in 4V ChP (*p-adj* < 0.05). Functions of the top ten upregulated and downregulated genes include solute transport, signaling, immune response, and trafficking (Table 2). While there is significant overlap in altered gene expression between the ChPs (Fig. 1A), the 4V ChP seems more susceptible to loss of *Otx2* activity. The response to *Otx2* knockdown results in a rather even distribution of upregulation (522 genes) and downregulation (392 genes) when grouping both 4V and LV ChPs. Interestingly, both ChPs have similar ontology enrichment in downregulated genes, indicated that *Otx2* is generally important for the expression of membrane proteins, glycoproteins, signaling proteins and cell adhesion proteins (Fig. 1B,C). While some of these functions are recapitulated in the upregulated genes, there is much more heterogeneity between the LV and 4V ChP. The LV ChP shows more immune response ontology while the 4V ChP shows more signaling related ontology. This suggests that conditional knockdown of *Otx2* leads to altered ChP barrier function and ChP signaling, and can impact immune responses.

**Table 1.**
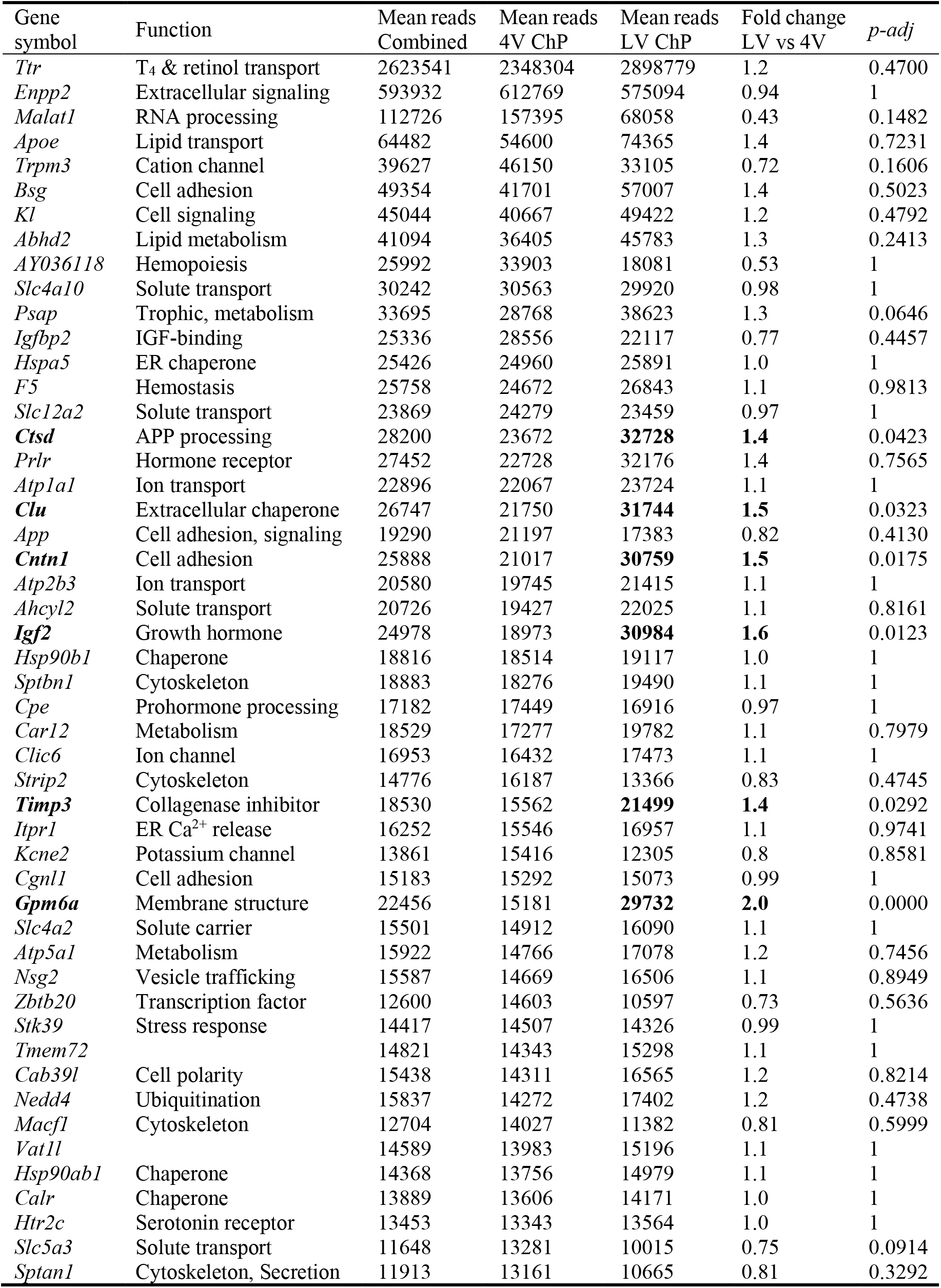
Top 50 genes expressed in lateral ventricle and 4^th^ ventricle choroid plexus. Significantly different expression between structures is indicated in bold.

| Gene symbol | Function | Mean reads Combined | Mean reads 4V ChP | Mean reads LV ChP | Fold change LV vs 4V | *p*-adj |
| --- | --- | --- | --- | --- | --- | --- |
| *Ttr* | T<sub>4</sub> & retinol transport | 2623541 | 2348304 | 2898779 | 1.2 | 0.4700 |
| *Enpp2* | Extracellular signaling | 593932 | 612769 | 575094 | 0.94 | 1 |
| *Malat1* | RNA processing | 112726 | 157395 | 68058 | 0.43 | 0.1482 |
| *Apoe* | Lipid transport | 64482 | 54600 | 74365 | 1.4 | 0.7231 |
| *Trpm3* | Cation channel | 39627 | 46150 | 33105 | 0.72 | 0.1606 |
| *Bsg* | Cell adhesion | 49354 | 41701 | 57007 | 1.4 | 0.5023 |
| *Kl* | Cell signaling | 45044 | 40667 | 49422 | 1.2 | 0.4792 |
| *Abhd2* | Lipid metabolism | 41094 | 36405 | 45783 | 1.3 | 0.2413 |
| *AY036118* | Hemopoiesis | 25992 | 33903 | 18081 | 0.53 | 1 |
| *Slc4a10* | Solute transport | 30242 | 30563 | 29920 | 0.98 | 1 |
| *Psap* | Trophic, metabolism | 33695 | 28768 | 38623 | 1.3 | 0.0646 |
| *Igfbp2* | IGF-binding | 25336 | 28556 | 22117 | 0.77 | 0.4457 |
| *Hspa5* | ER chaperone | 25426 | 24960 | 25891 | 1.0 | 1 |
| *F5* | Hemostasis | 25758 | 24672 | 26843 | 1.1 | 0.9813 |
| *Slc12a2* | Solute transport | 23869 | 24279 | 23459 | 0.97 | 1 |
| ***Ctsd*** | APP processing | 28200 | 23672 | **32728** | **1.4** | 0.0423 |
| *Prlr* | Hormone receptor | 27452 | 22728 | 32176 | 1.4 | 0.7565 |
| *Atp1a1* | Ion transport | 22896 | 22067 | 23724 | 1.1 | 1 |
| ***Clu*** | Extracellular chaperone | 26747 | 21750 | **31744** | **1.5** | 0.0323 |
| *App* | Cell adhesion, signaling | 19290 | 21197 | 17383 | 0.82 | 0.4130 |
| ***Cntn1*** | Cell adhesion | 25888 | 21017 | **30759** | **1.5** | 0.0175 |
| *Atp2b3* | Ion transport | 20580 | 19745 | 21415 | 1.1 | 1 |
| *Ahcyl2* | Solute transport | 20726 | 19427 | 22025 | 1.1 | 0.8161 |
| ***Igf2*** | Growth hormone | 24978 | 18973 | **30984** | **1.6** | 0.0123 |
| *Hsp90b1* | Chaperone | 18816 | 18514 | 19117 | 1.0 | 1 |
| *Sptbn1* | Cytoskeleton | 18883 | 18276 | 19490 | 1.1 | 1 |
| *Cpe* | Prohormone processing | 17182 | 17449 | 16916 | 0.97 | 1 |
| *Car12* | Metabolism | 18529 | 17277 | 19782 | 1.1 | 0.7979 |
| *Clic6* | Ion channel | 16953 | 16432 | 17473 | 1.1 | 1 |
| *Strip2* | Cytoskeleton | 14776 | 16187 | 13366 | 0.83 | 0.4745 |
| ***Timp3*** | Collagenase inhibitor | 18530 | 15562 | **21499** | **1.4** | 0.0292 |
| *Itpr1* | ER Ca<sup>2+</sup> release | 16252 | 15546 | 16957 | 1.1 | 0.9741 |
| *Kcne2* | Potassium channel | 13861 | 15416 | 12305 | 0.8 | 0.8581 |
| *Cgnl1* | Cell adhesion | 15183 | 15292 | 15073 | 0.99 | 1 |
| ***Gpm6a*** | Membrane structure | 22456 | 15181 | **29732** | **2.0** | 0.0000 |
| *Slc4a2* | Solute carrier | 15501 | 14912 | 16090 | 1.1 | 1 |
| *Atp5a1* | Metabolism | 15922 | 14766 | 17078 | 1.2 | 0.7456 |
| *Nsg2* | Vesicle trafficking | 15587 | 14669 | 16506 | 1.1 | 0.8949 |
| *Zbtb20* | Transcription factor | 12600 | 14603 | 10597 | 0.73 | 0.5636 |
| *Stk39* | Stress response | 14417 | 14507 | 14326 | 0.99 | 1 |
| *Tmem72* |  | 14821 | 14343 | 15298 | 1.1 | 1 |
| *Cab39l* | Cell polarity | 15438 | 14311 | 16565 | 1.2 | 0.8214 |
| *Nedd4* | Ubiquitination | 15837 | 14272 | 17402 | 1.2 | 0.4738 |
| *Macf1* | Cytoskeleton | 12704 | 14027 | 11382 | 0.81 | 0.5999 |
| *Vat1l* |  | 14589 | 13983 | 15196 | 1.1 | 1 |
| *Hsp90ab1* | Chaperone | 14368 | 13756 | 14979 | 1.1 | 1 |
| *Calr* | Chaperone | 13889 | 13606 | 14171 | 1.0 | 1 |
| *Htr2c* | Serotonin receptor | 13453 | 13343 | 13564 | 1.0 | 1 |
| *Slc5a3* | Solute transport | 11648 | 13281 | 10015 | 0.75 | 0.0914 |
| *Sptan1* | Cytoskeleton, Secretion | 11913 | 13161 | 10665 | 0.81 | 0.3292 |

**Table 2.** Top 10 differentially expressed genes in choroid plexus of Cre^+^ *Otx2^lox/lox^* mice.

| Gene symbol | Mean reads<br>vehicle | Mean reads<br>Cre-Tat | Fold<br>change | *p*-adj |
| --- | --- | --- | --- | --- |
| *Up-regulated in lateral ventricle ChP* |  |  |  |  |
| *Igkv1-135* | 0.6 | 33 | 61 | 0.0000 |
| *Slc1a6* | 2.1 | 82 | 39 | 0.0000 |
| *Mup5* | 123 | 4119 | 34 | 0.0000 |
| *Gpx3* | 295 | 7229 | 25 | 0.0000 |
| *Ighv1-67* | 1.6 | 36 | 22 | 0.0000 |
| *Saa3* | 2.2 | 47 | 22 | 0.0000 |
| *Tnfrsf11b* | 6.3 | 116 | 18 | 0.0000 |
| *Cacnb3* | 24 | 325 | 13 | 0.0000 |
| *Ndnf* | 89 | 1098 | 12 | 0.0000 |
| *Gm4841* | 3.8 | 44 | 12 | 0.0000 |
| *Down-regulated in lateral ventricle ChP* |  |  |  |  |
| *Ngfr* | 444 | 83 | -5.3 | 0.0000 |
| *Nrn1* | 1924 | 496 | -3.9 | 0.0000 |
| *B3galt2* | 109 | 31 | -3.6 | 0.0007 |
| *Dazl* | 207 | 60 | -3.4 | 0.0000 |
| *Itga10* | 184 | 54 | -3.4 | 0.0000 |
| *Slc26a7* | 1670 | 533 | -3.1 | 0.0000 |
| *Steap1* | 2224 | 735 | -3.0 | 0.0000 |
| *Defb11* | 473 | 169 | -2.8 | 0.0000 |
| *Entpd3* | 240 | 87 | -2.8 | 0.0001 |
| *Ccl9* | 2037 | 774 | -2.6 | 0.0000 |
| *Up-regulated in 4th ventricle ChP* |  |  |  |  |
| *9030619P08Rik* | 0.00 | 28.6 | infinite | 0.0000 |
| *Tmigd1* | 0.00 | 33.1 | infinite | 0.0000 |
| *A730020M07Rik* | 1.3 | 76 | 61 | 0.0000 |
| *Gpx3* | 181 | 10780 | 59 | 0.0000 |
| *Cacnb3* | 18 | 778 | 43 | 0.0000 |
| *Mup5* | 26 | 1053 | 41 | 0.0000 |
| *Fmod* | 286 | 8688 | 30 | 0.0000 |
| *Slitrk6* | 4.7 | 125 | 27 | 0.0000 |
| *Ndnf* | 84 | 2070 | 25 | 0.0000 |
| *Adcy8* | 10 | 233 | 23 | 0.0000 |
| *Down-regulated in 4th ventricle ChP* |  |  |  |  |
| *Ngfr* | 684 | 159 | -4.3 | 0.0000 |
| *Steap1* | 1225 | 326 | -3.8 | 0.0000 |
| *Elfn1* | 104 | 28 | -3.7 | 0.0000 |
| *Gnmt* | 122 | 33 | -3.7 | 0.0000 |
| *Gm22650* | 141 | 40 | -3.5 | 0.0000 |
| *Mir448* | 95 | 28 | -3.4 | 0.0002 |
| *Igf2os* | 89 | 26 | -3.4 | 0.0005 |
| *B3galt2* | 141 | 44 | -3.2 | 0.0000 |
| *Slc26a7* | 843 | 271 | -3.1 | 0.0000 |
| *Crhr2* | 504 | 164 | -3.1 | 0.0000 |

**Fig. 1.**
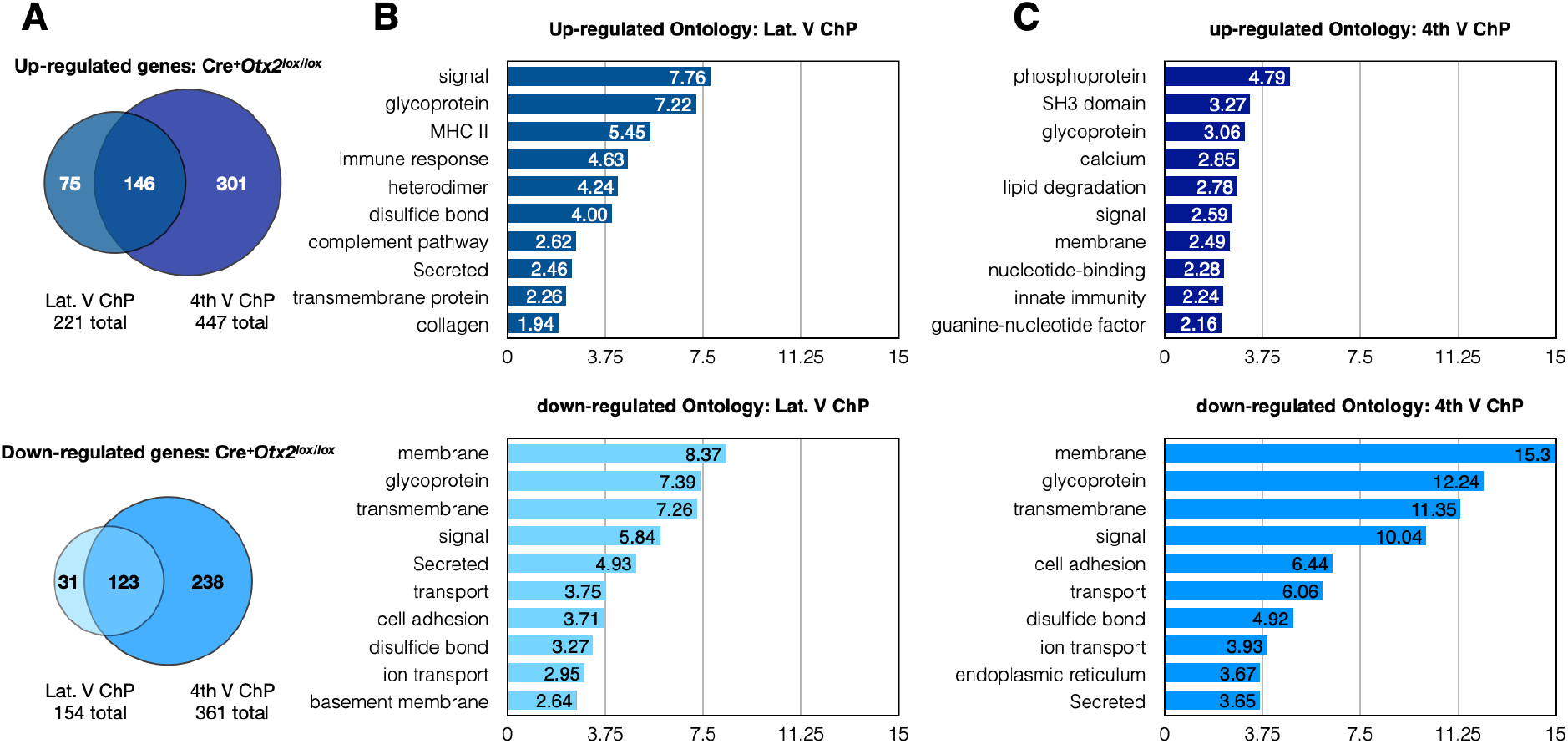
Changes in gene expression after *Otx2* knockdown in choroid plexus. (A) Venn diagrams of the number of up- or down-regulated genes (*p-adj* > 0.05) to compare overlap between lateral ventricle (LV ChP) and 4^th^ (B) ventricle (4V ChP). Ontology analysis of differentially regulated genes in lateral ventricle ChP. (C) Ontology analysis of differentially regulated genes in 4^th^ ventricle ChP.

The constitutive heterozygote *Otx2^+/GFP^* adult mice showed significant expression changes in 528 genes of the 4V ChP (*p-adj* < 0.05), which is comparatively less than for conditional *Otx2* knockdown in 4V ChP (Fig. 2A). Given that fewer genes are deregulated in this constitutive model, compensatory mechanisms for countering reduced OTX2 levels may have been activated during development. The changes in gene expression were relatively balanced between upregulation (273 genes) and downregulation (255 genes), and ontology analysis revealed shared terms including glycoprotein, signal, membrane-related, and secreted proteins (Fig. 2B). The upregulated genes are also enriched for cell adhesion and alternative splicing, while downregulated genes are enriched for trafficking and transport. This suggests that brain-wide and life-long knockdown of *Otx2* leads to altered ChP signaling, barrier functions, and brain homeostasis.

**Fig. 2.**
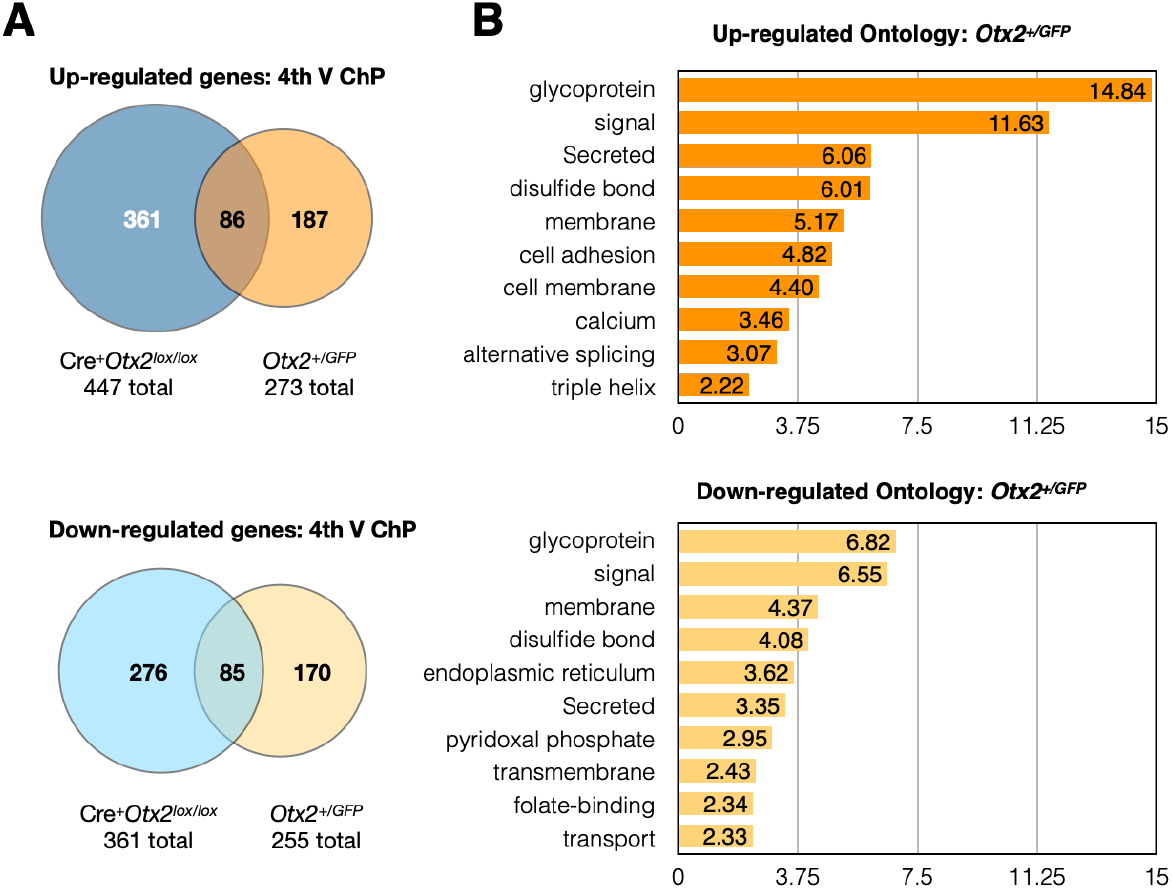
Gene expression in choroid plexus of *Otx2^+/GFP^* mice. (A) Venn diagrams of the number of up- or down-regulated genes (*p-adj* > 0.05) to compare overlap between 4^th^ ventricle (4V ChP) changes in *Otx2^+/GFP^* mice and in Cre-Tat icv-injected *Otx2^lox/lox^* mice. (B) Ontology analysis of differentially regulated genes in *Otx2^+/GFP^* mice.

We hypothesized that genes deregulated in both conditional and constitutive models could be either direct targets of OTX2 transcription regulation or targets of important OTX2-dependent pathways. Comparison of gene expression changes in 4V ChP of these two models revealed an overlap of more than 80 genes in both upregulated or downregulated repertoires (Fig. 2A). This represented about half the identified expression changes in *Otx2^+/GFP^* mice, but less than a third of changes in the conditional model. When genes from LV ChP conditional *Otx2* knockdown are included in the analysis, we identified 42 genes globally upregulated and 34 genes globally downregulated (Table 3). To determine whether this list contains direct OTX2 transcription targets, we compared it with OTX2 chromatin-immunoprecipitation experiments previously performed in mouse embryonic brain (Hoch et al., 2015) or adult retina (Samuel et al., 2014). However, we find almost no overlap, with only *Ttr* being a common target. This suggests that transcriptional activity of OTX2 has different targets in the adult choroid plexus and/or that these deregulated genes are downstream targets of OTX2-dependent pathways. It will be necessary to perform ChIPseq analysis of adult choroid plexus to distinguish between these possibilities. Taken together, our analysis identifies new potential functions for *Otx2* in the adult brain. We find upregulation of immune factors, specifically in the conditional *Otx2* loss-of-function model, and deregulation of genes involved in cellular adhesion, trafficking, signaling, and secretion in both knockdown models, suggesting altered ChP function and disruption of ChP barriers.

**Table 3.**
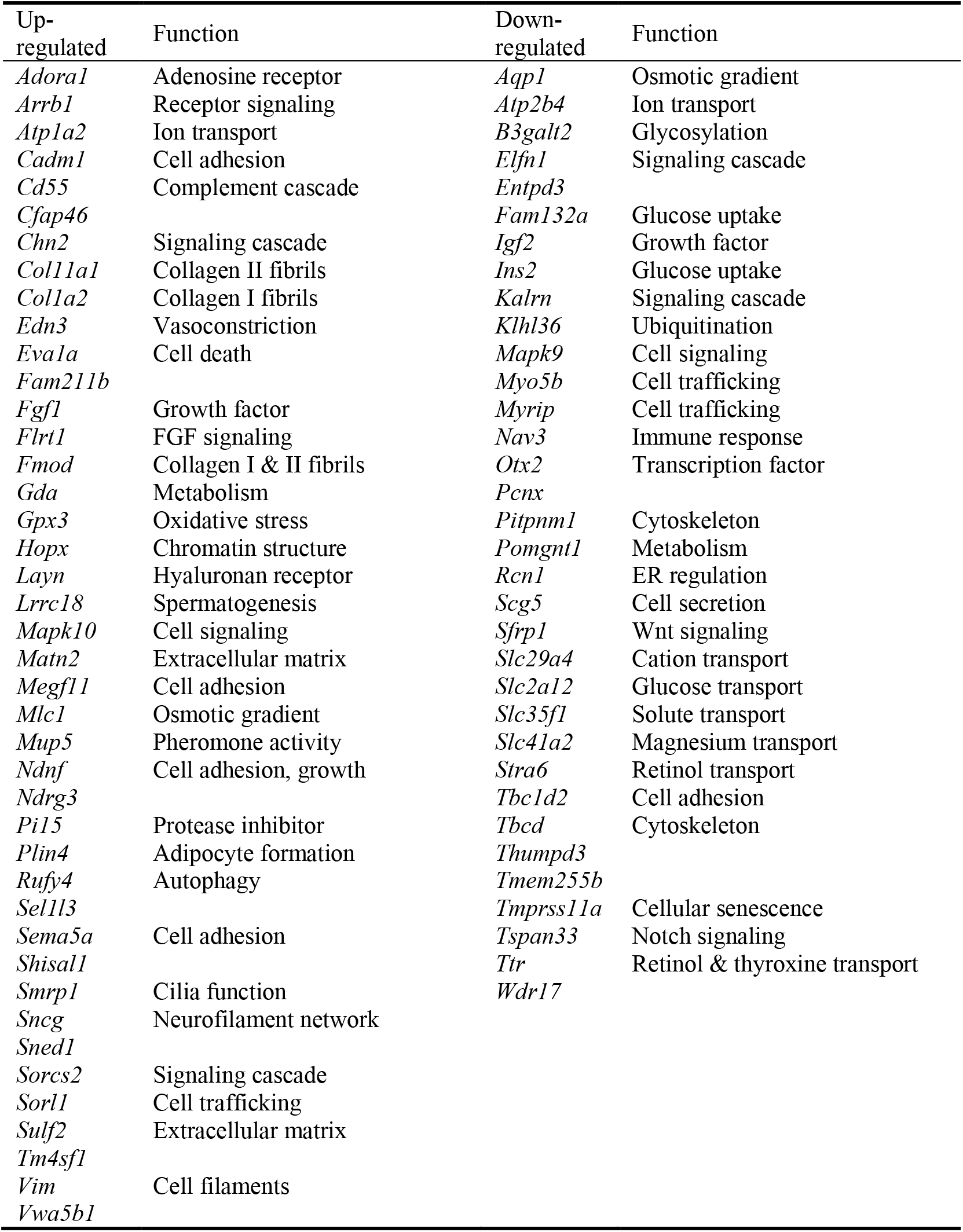
Choroid plexus genes with significant expression changes in both *Otx2* knockdown experiments, including lateral ventricle and 4^th^ ventricle from Cre^+^*Otx2^lox/lox^* mice and 4^th^ ventricle from *Otx2^+/GFP^* mice.

### Altered expression of ChP secreted factors

Given that our various ontology analyses often evoked secreted factors, we focused on ChP factors secreted in CSF and implicated in embryonic and/or adult neurogenesis (Table 4), which is one of the recently identified functions of adult ChP (Falcão et al., 2012; Silva-Vargas et al., 2016). Factors implicated in embryonic neurogenesis include SHH, BMPs and WNTs (Bjornsson et al., 2015). While *Shh* expression was not observed (mean reads <1) in either ChP of WT mice, as previously reported (Lun et al., 2015), only *Otx2^+/GFP^* mice (and not conditional *Otx2* ChP knockdown mice) showed significant increase in *Shh* expression. Between the various *Bmp* and *Wnt* family genes, only *Bmp7* and *Wnt2b* were differentially expressed in Cre^+^*Otx2^lox/lox^* mice as compared to Veh^+^*Otx2^lox/lox^* mice. Canonical Wnt signaling is perturbed in embryos with conditional *Otx2* ChP knockdown, and it has been proposed that WNT4 and TGM2, enriched in the CSF of mutant mice, are the effectors (Johansson et al., 2013). While *Wnt4* was poorly expressed (~20 mean reads) and unchanged in both LV and 4V ChP of Cre^+^*Otx2^lox/lox^* mice, *Tgm2* was significantly upregulated in 4V ChP, suggesting OTX2 retains some embryonic functions in the adult, such as repressing *Tgm2* expression. Given that *Otx2* knockdown experiments in late embryonic development showed OTX2 is necessary for 4V but not LV ChP maintenance (Johansson et al., 2013), these results suggest that adult *Otx2* expression could retain this maintenance function in 4V ChP.

**Table 4.**
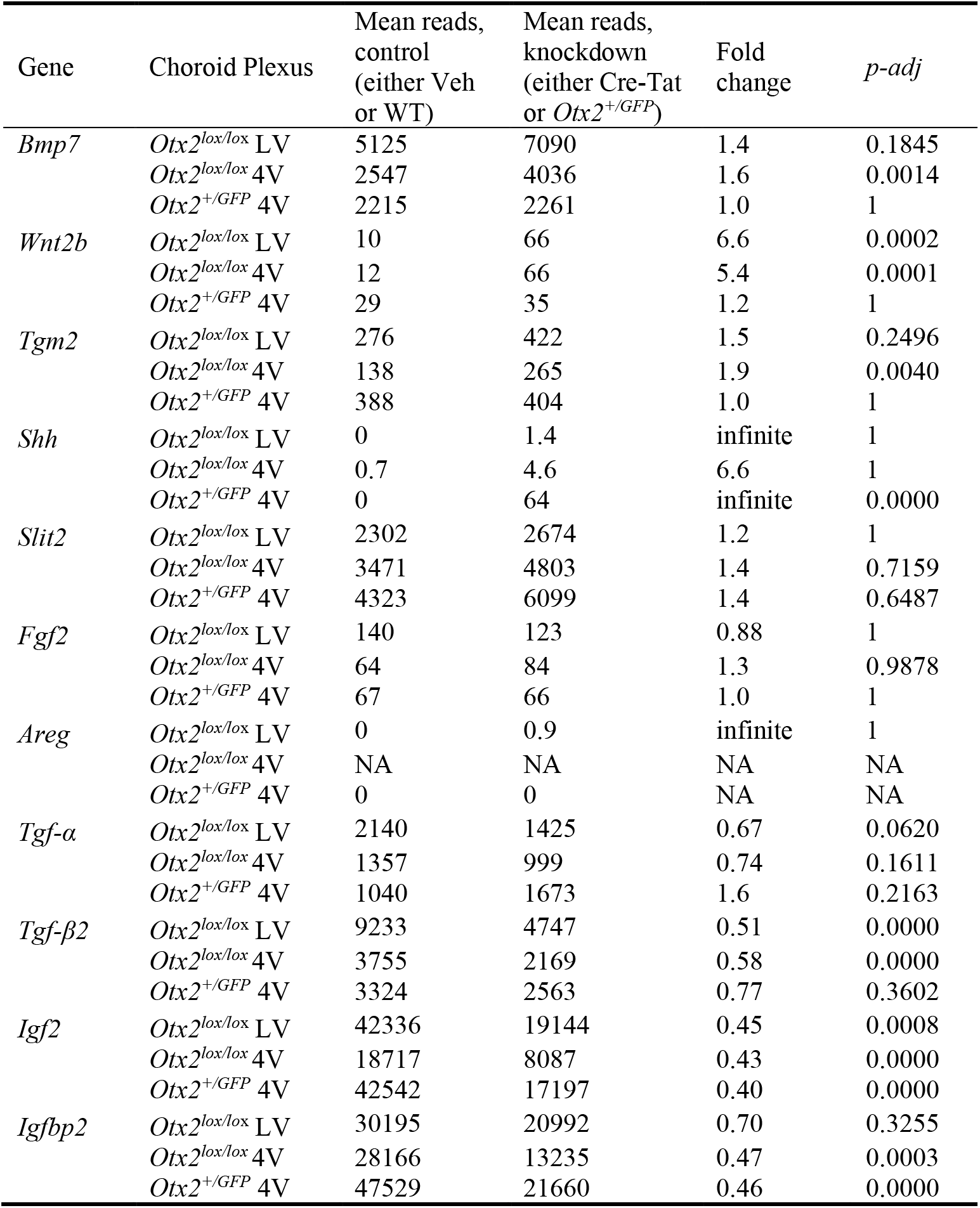
Expression of secreted factors in *Otx2* knockdown experiments.

| Gene | Choroid Plexus | Mean reads,<br>control<br>(either Veh<br>or WT) | Mean reads,<br>knockdown<br>(either Cre-Tat<br>or *Otx2*<sup>+/GFP</sup>) | Fold<br>change | *p*-adj |
| --- | --- | --- | --- | --- | --- |
| *Bmp7* | *Otx2*<sup>lox/lox</sup> LV | 5125 | 7090 | 1.4 | 0.1845 |
|  | *Otx2*<sup>lox/lox</sup> 4V | 2547 | 4036 | 1.6 | 0.0014 |
|  | *Otx2*<sup>+/GFP</sup> 4V | 2215 | 2261 | 1.0 | 1 |
| *Wnt2b* | *Otx2*<sup>lox/lox</sup> LV | 10 | 66 | 6.6 | 0.0002 |
|  | *Otx2*<sup>lox/lox</sup> 4V | 12 | 66 | 5.4 | 0.0001 |
|  | *Otx2*<sup>+/GFP</sup> 4V | 29 | 35 | 1.2 | 1 |
| *Tgm2* | *Otx2*<sup>lox/lox</sup> LV | 276 | 422 | 1.5 | 0.2496 |
|  | *Otx2*<sup>lox/lox</sup> 4V | 138 | 265 | 1.9 | 0.0040 |
|  | *Otx2*<sup>+/GFP</sup> 4V | 388 | 404 | 1.0 | 1 |
| *Shh* | *Otx2*<sup>lox/lox</sup> LV | 0 | 1.4 | infinite | 1 |
|  | *Otx2*<sup>lox/lox</sup> 4V | 0.7 | 4.6 | 6.6 | 1 |
|  | *Otx2*<sup>+/GFP</sup> 4V | 0 | 64 | infinite | 0.0000 |
| *Slit2* | *Otx2*<sup>lox/lox</sup> LV | 2302 | 2674 | 1.2 | 1 |
|  | *Otx2*<sup>lox/lox</sup> 4V | 3471 | 4803 | 1.4 | 0.7159 |
|  | *Otx2*<sup>+/GFP</sup> 4V | 4323 | 6099 | 1.4 | 0.6487 |
| *Fgf2* | *Otx2*<sup>lox/lox</sup> LV | 140 | 123 | 0.88 | 1 |
|  | *Otx2*<sup>lox/lox</sup> 4V | 64 | 84 | 1.3 | 0.9878 |
|  | *Otx2*<sup>+/GFP</sup> 4V | 67 | 66 | 1.0 | 1 |
| *Areg* | *Otx2*<sup>lox/lox</sup> LV | 0 | 0.9 | infinite | 1 |
|  | *Otx2*<sup>lox/lox</sup> 4V | NA | NA | NA | NA |
|  | *Otx2*<sup>+/GFP</sup> 4V | 0 | 0 | NA | NA |
| *Tgf-α* | *Otx2*<sup>lox/lox</sup> LV | 2140 | 1425 | 0.67 | 0.0620 |
|  | *Otx2*<sup>lox/lox</sup> 4V | 1357 | 999 | 0.74 | 0.1611 |
|  | *Otx2*<sup>+/GFP</sup> 4V | 1040 | 1673 | 1.6 | 0.2163 |
| *Tgf-β2* | *Otx2*<sup>lox/lox</sup> LV | 9233 | 4747 | 0.51 | 0.0000 |
|  | *Otx2*<sup>lox/lox</sup> 4V | 3755 | 2169 | 0.58 | 0.0000 |
|  | *Otx2*<sup>+/GFP</sup> 4V | 3324 | 2563 | 0.77 | 0.3602 |
| *Igf2* | *Otx2*<sup>lox/lox</sup> LV | 42336 | 19144 | 0.45 | 0.0008 |
|  | *Otx2*<sup>lox/lox</sup> 4V | 18717 | 8087 | 0.43 | 0.0000 |
|  | *Otx2*<sup>+/GFP</sup> 4V | 42542 | 17197 | 0.40 | 0.0000 |
| *Igfbp2* | *Otx2*<sup>lox/lox</sup> LV | 30195 | 20992 | 0.70 | 0.3255 |
|  | *Otx2*<sup>lox/lox</sup> 4V | 28166 | 13235 | 0.47 | 0.0003 |
|  | *Otx2*<sup>+/GFP</sup> 4V | 47529 | 21660 | 0.46 | 0.0000 |

We have previously shown that OTX2 secreted into the CSF from the ChP can regulate adult neurogenesis non-cell autonomously by transferring into astrocytes in the V-SVZ and RMS, thereby affecting neuroblast migration (Planques et al., 2019). This study also showed that ChP *Otx2* knockdown in Cre^+^*Otx2^lox/lox^* adult mice, which will impact both cell- and non-cell-autonomous activity, also led to significantly reduced neurogenesis, suggesting that cell-autonomous effects on ChP targets may also be involved. IGF2 and SLIT1/2 have been found to regulate both embryonic and adult neurogenesis (Lehtinen et al., 2011; Nguyen-Ba-Charvet et al., 2004; Sawamoto et al., 2006; Ziegler et al., 2012). While *Slit1* is not expressed and *Slit3* is only weakly expressed in adult ChP, *Slit2* is highly expressed but shows no significant change in expression in ChP with reduced *Otx2*. Admittedly, there is a trend towards increased *Slit2* expression and we cannot exclude the potential for biologically relevance. *Igf2* was significantly downregulated more than 2-fold in all ChPs upon *Otx2* knockdown. However, there was a concomitant downregulation in *Igfbp2*, which can inhibit IGF2, suggesting the level of IGF2 activity could be maintained through compensatory reduction in inhibiting factors. Other factors influencing adult neurogenesis include amphiregulin (AREG) (Falk and Frisén, 2002), FGF2 (Douet et al., 2013; Hayamizu et al., 2001; Jin et al., 2003), and TGF-α (Tropepe et al., 1997), yet we found no significant change in their expression (and no detectible expression of *Areg*). Finally, other factors show more change in gene expression after acute *Otx2* knockdown compared to constitutive knockdown. TGF-β negatively regulates adult neurogenesis (Wachs et al., 2006), and *Tgf-β2* is downregulated in both ChP of Cre^+^*Otx2^lox/lox^* mice (Table 4). Taken together, these minimal or compensatory changes in specific secreted signaling factors suggest that cell-autonomous OTX2 activity in ChP could have only a minor role in regulating adult neurogenesis. This hypothesis is consistent with previously reported effects of ChPl OTX2 knockdown; decreased neurogenesis was mainly attributed to the reduction of OTX2 CSF levels and its non-cell-autonomous activity in V-SVZ and RMS (Planques et al., 2019).

### Altered expression of immune and stress factors

Given the altered expression of homeostasis and stress-response related factors in both ChPs of conditional *Otx2* knockdown mice, we turned to viral expression of shRNA-*Otx2* in LV and 4V ChPs to achieve greater reduction in *Otx2* expression. The use of intracerebroventricular-injected AAV5 provides for ChP-specific expression (Arnaud et al., 2019; Watson et al., 2005). Through qPCR analysis, we measured a 70% decrease in (mRNA) *Otx2* and a concomitant very large decrease in expression of a known direct transcriptional target, transthyretin (*Ttr*) (Fig. 3A). TTR, the most highly expressed protein in ChP (Table 1), is secreted into CSF and transports thyroxin and retinol-binding protein, and has a role in cognition and memory, psychological health and emotion (for recent review, see (Sharma et al., 2019)). Given that OTX2 directly regulates *Ttr* expression, this strongly implies that *Otx2* expression levels in ChP impact brain function. Furthermore, the downregulation of aquaporin (*Aqp)* (Table 3) was also confirmed, specifically *Aqp1* and not *Aqp4* (Fig. 3A), suggesting OTX2 helps regulate CSF water homeostasis.

**Fig. 3.**
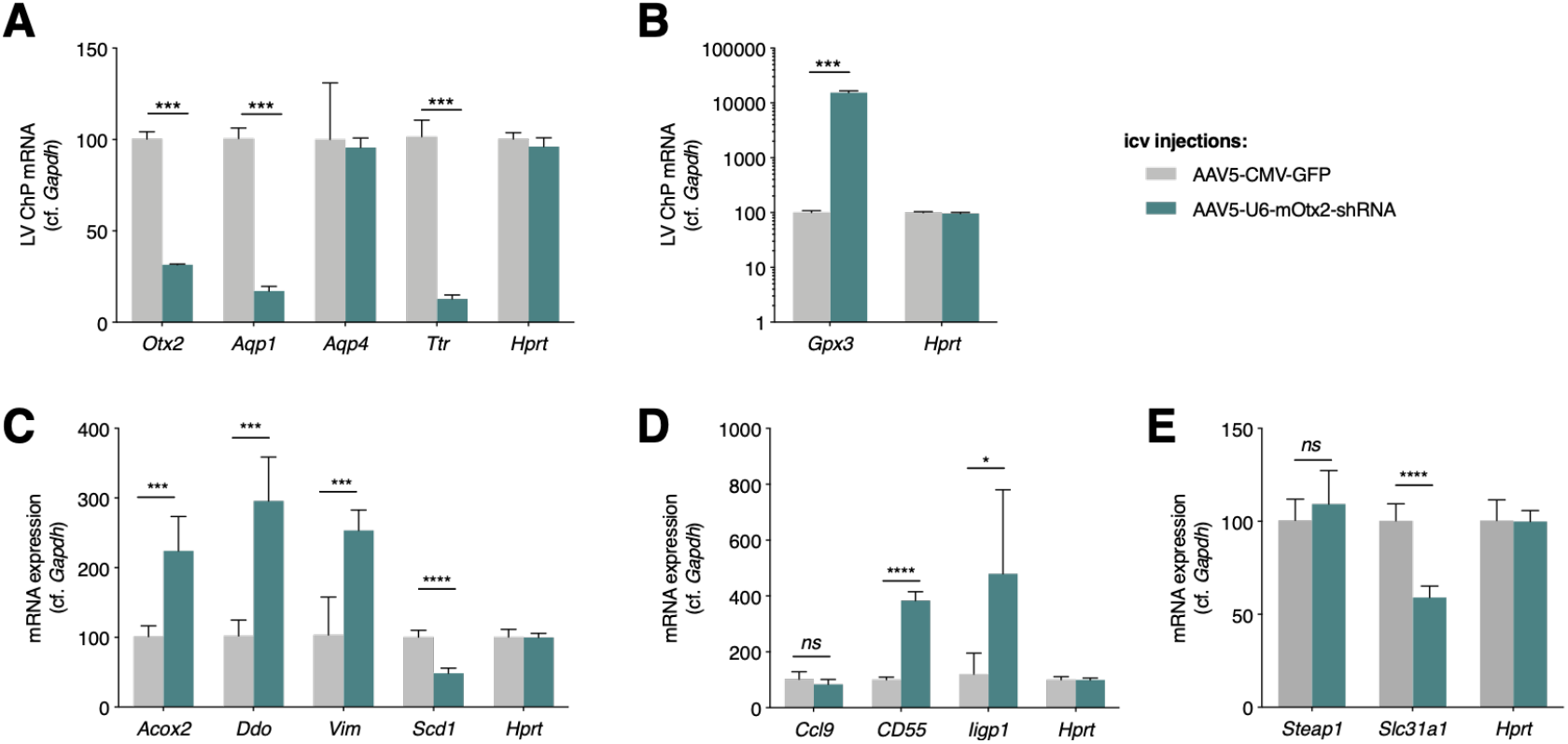
OTX2 regulates choroid plexus expression of oxidative stress, immune system, and metal transport genes. Quantitative PCR analysis of lateral ventricle (LV) ChP gene expression in wildtype mice after viral expression of shRNA against mouse (mRNA) *Otx2* (shRNA-*Otx2*). (A) Analysis of control genes to validate shRNA-*Otx2* activity. (B) *Otx2* knockdown induces high expression of *Gpx3*. (C) Analysis of select genes involved in oxidative stress response. (D) Analysis of select genes involved in immune system response. (E) Analysis of select genes involved in metal ion transport.

In keeping with roles in brain homeostasis and surveillance, we also chose targets from ontology analysis (Fig. 1) with functions related to oxidative stress, immune response and metal ion transport. A surprising finding was the over 100-fold increase in glutathione peroxidase 3 (*Gpx3*), an extracellular enzyme that catalyzes the reduction of peroxidases and protects cells from oxidative damage, suggesting loss of OTX2 has a significant impact on cell physiology (Fig. 3B). Other compensatory mechanisms against oxidative stress include decreased fatty-acid oxidation (*Scd1*), increased peroxisome function (*Acox2*, *Ddo*) for countering oxidative stress and inflammation (Terlecky, 2012), and changes in structural cell response (*Vim*) (De Pablo et al., 2013) (Fig. 3C). In relation to immune response (Fig. 3D), we verified a complement activation factor (*CD55*), an inflammatory response chemokine (*Ccl9*), and an innate immune response factor (*Iigp1*). The direction of change in expression of all of these factors upon acute viral *Otx2* knockdown was consistent with the constitutive and conditional mouse models. It remains unclear whether loss of OTX2 provokes oxidative stress, and thus indirect activation of genes such as *Gpx3*, or whether OTX2 regulates genes involved in reactive oxygen species signaling and/or stress response. Finally, we also quantified factors related to brain homeostasis and metal ion transport (*Steap1* and *Slc31a*) with altered expression in the conditional Cre^+^*Otx2^lox/lox^* mice, (Fig. 3E). Only *Slc31a*, which transports copper ions, had concomitant reduced expression upon *Otx2* knockdown. Together these findings suggest that ChP function is greatly impacted by *Otx2* expression level, opening the question whether *Otx2* overexpression in ChP would also deregulate homeostasis and elicit immune responses in a wildtype context or on the contrary rescue deficits in homeostasis in an aged or diseased animal.

### Otx2 protein interactions

To further analyze OTX2 function in adult ChP, we performed several OTX2 co-immunoprecipitation (co-IP) experiments with mass spectrometry (MS) analysis to identify potential protein partners. We previously discovered that OTX2 protein is secreted by the ChP into CSF and accumulates non-cell autonomously in V-SVZ and RMS astrocytes (Planques et al., 2019) and in VCx parvalbumin cells (Kim et al., 2014; Spatazza et al., 2013). The identification of alternate protein partners in cell-autonomous and non-cell-autonomous contexts would suggest OTX2 takes on specific roles after transferring between cells. In order to test this hypothesis, and to reinforce ChP analysis, we also performed OTX2 co-IP on lysates from adult mouse V-SVZ, RMS and VCx.

We used three criteria to obtain a list of potential OTX2 protein interactions in the four brain structures (Table 5): (i) unique proteins with 3 or more peptides identified exclusively in OTX2 compared to IgG co-IP samples (Unique protein, ≥ 3 peptides); (ii) proteins identified with 3 or more peptides in OTX2 co-IP samples and having a relative peptide difference greater than 50% compared to IgG co-IP (Selected protein, ≥ 50% rel. ∆); and (iii) all small proteins (≤ 25 kDa) exclusive to OTX2 co-IP samples regardless of peptide number (Unique small protein, ≤ 25 kDa) given that they have fewer identifiable MS peptides. These lists were used for comparison between structures and ontology analysis. We generated a list of 60 high-confidence protein partners of OTX2 in ChP that were common to all three ChP samples (Table 6). These proteins cover a limited range of functions, including cell adhesion, cell trafficking, cell signaling, metabolism, RNA binding, RNA processing, transcription, chromatin structure and DNA repair. Interestingly, more than 10% (8 proteins) belong to the “tier 1” proteins identified in stress granules (Youn et al., 2019), which are involved in translational control and post-transcriptional regulation. Although this functional class was not identified by KEGG pathway analysis (see below), this is likely due to the absence of annotation given the only recent emergence of updated comprehensive inventories of stress granule proteins. Thus, we can only hypothesize that OTX2 interacts with these granules, although this putative function is given weight by the presence of the PAX1 homeoprotein among the “tier 1” proteins, by the *in vivo* interaction between EMX2 homeoprotein with translation initiation factor eIF4E (Nédélec et al., 2004), and by the involvement of PROX1 homeoprotein in liquid-liquid phase separation (Liu et al., 2020), which also underlies stress granule assembly (Youn et al., 2019). Also of note are the putative partners MECP2 and MOV10, given that OTX2 impacts MECP2 foci in the postnatal mouse visual and auditory cortex (Apulei et al., 2019), and that both OTX2 and EN1 homeoproteins may regulate transposable elements (Blaudin de Thé et al., 2018; Guo et al., 2021).

**Table 5.** Summary of protein lists and criteria for identification of putative OTX2 protein partners.

| List name | Choroid<br>plexus | V-SVZ | RMS | Visual<br>cortex |
| --- | --- | --- | --- | --- |
| Total proteins OTX2 | 4814 | 1138 | 2425 | 2644 |
| Total proteins IgG | 3602 | 1776 | 2274 | 2667 |
| Unique proteins OTX2 ($\geq 3$ peptides) | 392 | 17 | 40 | 29 |
| Unique proteins IgG ($\geq 3$ peptides) | 59 | 139 | 22 | 25 |
| Selected proteins OTX2 ($\geq 50\%$ rel. $\Delta$) | 653 | 6 | 75 | 37 |
| Unique small proteins Otx2 ($\leq 25$ kDa) | 182 | 31 | 68 | 48 |
| Total OTX2 partners | 1195 of 4814 | 52 of 1138 | 180 of 2425 | 109 of 2644 |

**Table 6.** High confidence OTX2 putative protein partners in choroid plexus identified by MS analysis. Proteins indicated in bold are unique for OTX2 co-IP, proteins with * are “tier 1” stress granule proteins.

| Protein | Function | Protein | Function |
| --- | --- | --- | --- |
| ABCF1* | Translation | MAP4 | Cytoskeleton |
| ACOT11 | Lipid metabolism | MCM3AP | RNA export |
| AGO1* | RNA silencing | MECP2 | Transcription, epigenetics |
| APC | Cell adhesion | MLYCD | Metabolism |
| **ARHGEF6** | Trafficking | MOV10* | RNA and LINE-1 silencing |
| ARHGEF7 | Trafficking, Cell adhesion | MSI2* | Translation |
| ARVCF | Cell adhesion | MYCBP2 | Transcription |
| CDH2 | Cell adhesion | **NFATC2** | Signaling |
| CDH3 | Cell adhesion | PIKFYVE | Trafficking |
| CHD4 | Cell adhesion | PITPNM2 | Trafficking |
| CTNNA1 | Cell adhesion | POLDIP3 | Translation |
| CTNNA2 | Cell adhesion | PRRC2A* | RNA splicing, Stress granule |
| CTNNB1 | Cell adhesion | RBM39 | RNA splicing |
| DDX41 | RNA splicing | RHOT1 | Mitochondrial trafficking |
| EDC4* | RNA processing | RPL19 |  |
| EPB41L5 | Cell adhesion | RPL21 | Translation |
| ERBIN | Signaling | RPL22 |  |
| **FIG4** | Trafficking | RPL29 | Translation |
| FMNL3 | Cytoskeleton | RPL35 | Translation |
| GIT1 | Trafficking, Cell adhesion | RPL36A |  |
| **GIT2** | Trafficking | SRRM2 | RNA splicing |
| GJA1 | Gap junction | STRAP* | Stress response |
| GPAM | Metabolism | TJP2 | Cell adhesion |
| GTPBP1 | RNA processing | TMPO | Nuclear membrane |
| HNRPLL* | RNA splicing | TNS2 | Signaling |
| ILF2 | Transcription | TRPV4 | Osmotic sensitivity |
| KIFAP3 | Chromosome structure | **VAC14** | Trafficking |
| LBR | Metabolism | VRK3 | Signaling |
| LIG3 | DNA repair | **WDR70** |  |
| MAP1A | Cytoskeleton | ZFR | RNA export |

Few OTX2 partners have been biochemically and functionally validated. One key partner during embryogenesis is MEIS2, which is a co-activator for mesencephalon specification (Agoston and Schulte, 2009). *Meis2* is expressed in ChP at low levels (170 mean reads in 4V ChP; 31 mean reads in LV ChP) as compared to OTX2 (5954 mean reads in 4V ChP and 6829 mean reads in LV ChP) and appears not to be a major partner of OTX2 in ChP. TLE4 is another identified protein partner of OTX2 during development and allows repression of mesencephalon fate (Heimbucher et al., 2007). Despite its expression in ChP (798 mean reads in 4V ChP and 689 mean reads in LV ChP), TLE4 was not identified in our OTX2 co-IP, although this could be due to TLE4 being under the limit of detection or having peptides too hydrophobic or hydrophilic for MS detection. The potential absence of TLE4 suggests OTX2 protein interactions depend strongly on cell type and developmental context.

To identify novel OTX2 protein partners ubiquitous throughout the brain, we compared the lists from the four brain structures: ChP (pooled LV and 4V), V-SVZ, RMS and VCx. Few high-confidence proteins (Selected protein ≥ 50% rel. ∆) were common to the three non-cell autonomous structures (Table 7). Of these 14 common proteins, 5 were also identified in ChP. Interestingly, these top-ranked proteins include FIG4, VAC14 and PIKFYVE, which play a role in phosphatidylinositol(3,5)bisphosphate [PI(3,5)P2] regulation, in multivesicular body (MVB) biogenesis, and in endosome autophagy and trafficking (McCartney et al., 2014), suggesting OTX2 plays a role in vesicle transport or is carried via MVBs. Given that MVBs can give rise to extracellular vesicles, interaction with OTX2 may reinforce its role in regulating pathways of extracellular protein expression identified in our RNA sequencing analysis.

**Table 7.** Peptide number comparisons of OTX2 putative protein partners common to non-cell autonomous structures. Cell-autonomous partners are indicated in bold.

| Protein | Function | V-SVZ |  | RMS |  | VCx |  | ChP |  |
| --- | --- | --- | --- | --- | --- | --- | --- | --- | --- |
|  |  | Otx2 co-IP | IgG co-IP | Otx2 co-IP | IgG co-IP | Otx2 co-IP | IgG co-IP | Otx2 co-IP | IgG co-IP |
| ACIN1 | mRNA splicing | 3 | 0 | 11 | 1 | 16 | 4 |  |  |
| **ACOT11** | Lipid metabolism | 4 | 0 | 5 | 0 | 8 | 1 | **34** | **7** |
| ARCN1 | Protein transport | 3 | 1 | 7 | 2 | 4 | 1 |  |  |
| DDX46 | mRNA splicing | 30 | 0 | 45 | 1 | 34 | 3 |  |  |
| EIF4A3 | mRNA translation | 4 | 0 | 10 | 2 | 11 | 5 |  |  |
| **FIG4** | PI(3,5)P2 regulation, MVB | 20 | 0 | 26 | 0 | 27 | 0 | **43** | **0** |
| **KCND3** | Potassium channel | 6 | 0 | 5 | 0 | 7 | 0 | **6** | **0** |
| **PIKFYVE** | PI(3,5)P2 regulation, MVB | 36 | 0 | 42 | 0 | 59 | 0 | **122** | **1** |
| RBM25 | mRNA splicing | 5 | 0 | 11 | 1 | 12 | 3 |  |  |
| SF3A1 | mRNA splicing | 3 | 0 | 11 | 3 | 13 | 6 |  |  |
| SF3B1 | mRNA splicing | 3 | 0 | 22 | 0 | 25 | 9 |  |  |
| SNRNP200 | mRNA splicing | 5 | 0 | 14 | 1 | 31 | 4 |  |  |
| THOC2 | mRNA export | 3 | 0 | 5 | 0 | 3 | 0 |  |  |
| **VAC14** | PI(3,5)P2 regulation, MVB | 39 | 2 | 49 | 3 | 56 | 0 | **75** | **0** |

**Fig. 4.**
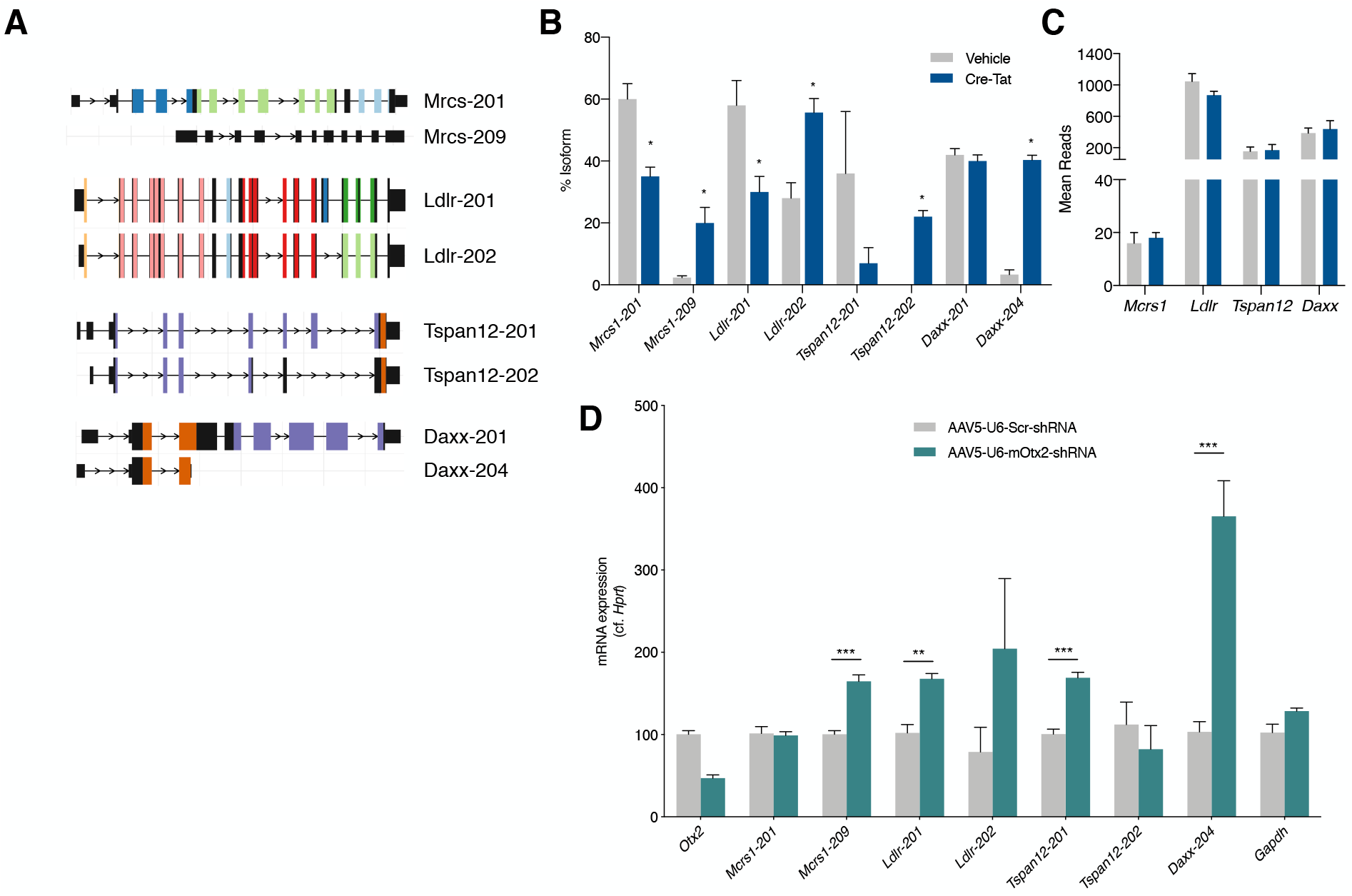
Analysis of splice variations induced by *Otx2* knockdown in choroid plexus. (A) Selected isoforms of genes of interest. Locus length is in arbitrary units. Colors represent different protein domains within a given gene. (B) Mean reads from transcriptomic analysis of lateral ventricle (LV) ChP from vehicle and Cre-Tat icv-injected *Otx2^lox/lox^* mice. (C) Isoform usage, shown as % of total isoforms, in LV ChP from vehicle and Cre-Tat icv-injected *Otx2^lox/lox^* mice. (D) Quantitative PCR analysis of LV ChP isoform expression in WT mice after viral expression of shRNA against mouse (mRNA) *Otx2* (shRNA-*Otx2*).

In order to identify potential differences between cell autonomous and non-cell autonomous partners, we performed KEGG pathway analysis on OTX2 protein partners for all structures individually (Table 8). No dramatic differences were found between structures, suggesting conserved roles of OTX2 in cell autonomous and non-cell autonomous OTX2 target structures. Common to nearly all structures are metabolic pathways, RNA transport, oxidative phosphorylation, RNA processing, and spliceosome. Pathways specific to ChP pertain to the maintenance of tight junctions, protein processing and actin cytoskeleton regulation. The enrichment of tight junction class was also identified in conditional Cre^+^*Otx2^lox/lox^* mice, suggesting involvement of OTX2 both in gene regulation and cellular functions for cell-autonomous ChP maintenance. Of the 14 proteins in common between VCx, V-SVZ and RMS, 8 of them are involved in RNA processing, suggesting a novel function for OTX2. Although the splicesome pathway was also enriched in ChP, these proteins stand out for their involvement in U5 snRNP complex, exon junction complex or mRNA export complex, whereas in the ChP the spliceosome proteins are either splicing co-factors, part of the SMN complex, or part of the U2 snRNP complex. Interestingly, OTX2 has been shown to bind initiation factor eIF4e in GST pull-down experiments (Nédélec et al., 2004), while other homeoproteins have been shown to bind translation machinery (Nédélec et al., 2004; Rezsohazy, 2014; Topisirovic et al., 2005) implicated in RNA export, transport, and translation. Taking the high-confidence partners together with KEGG pathway analysis, cell-autonomous OTX2 is likely implicated in the regulation of genomic landscape, the regulation and processing of RNA, the trafficking of signals, and the maintenance of cellular adhesion, while non-cell-autonomous OTX2 is more implicated in the processing of RNA.

**Table 8.** List of KEGG pathways associated with putative cell- and non-cell autonomous OTX2 protein partners.

| KEGG pathway | Size | adj P value |
| --- | --- | --- |
| *Choroid plexus (1326 proteins)* |  |  |
| Tight junction | 27 of 137 | 1.61E-15 |
| Metabolic pathways | 77 of 1184 | 3.97E-14 |
| Protein processing in ER | 26 of 169 | 1.10E-12 |
| RNA transport | 24 of 168 | 2.95E-11 |
| Ribosome biogenesis | 18 of 86 | 2.95E-11 |
| Regulation of actin cytoskeleton | 27 of 216 | 2.95E-11 |
| Ribosome | 19 of 119 | 7.41E-10 |
| Spliceosome | 20 of 138 | 1.22E-09 |
| Oxidative phosphorylation | 20 of 147 | 3.47E-09 |
| Axon guidance | 18 of 131 | 2.10E-08 |
| *V-SVZ (79 proteins)* |  |  |
| Ribosome | 15 of 119 | 1.44E-24 |
| Spliceosome | 15 of 138 | 7.45E-24 |
| RNA transport | 4 of 168 | 4.00E-04 |
| mRNA surveillance | 2 of 93 | 2.19E-02 |
| Neurotrophin signalling | 2 of 131 | 2.97E-02 |
| *RMS (219 proteins)* |  |  |
| Spliceosome | 28 of 138 | 1.38E-38 |
| Oxidative phosphorylation | 11 of 147 | 3.62E-10 |
| Metabolic pathways | 24 of 1184 | 6.10E-10 |
| Alzheimer disease | 10 of 188 | 3.51E-08 |
| mRNA surveillance | 8 of 93 | 3.51E-08 |
| *Visual cortex (192 proteins)* |  |  |
| Spliceosome | 17 of 138 | 2.06E-20 |
| RNA transport | 7 of 168 | 2.92E-05 |
| Metabolic pathways | 14 of 1184 | 5.00E-04 |
| Insulin signaling | 5 of 137 | 7.00E-04 |
| Oxidative phosphorylation | 5 of 147 | 7.00E-04 |

### Splice variant analysis

Given the high confidence of OTX2 interaction with spliceosome pathway proteins, we extended the analysis of our transcriptomic data of LV ChP from Cre^+^*Otx2^lox/lox^* mice to measure changes in splice variants. Isoform usage was found to be significantly changed in the coding transcripts for only 4 genes (*Mcrs*, *Ldlr*, *Tspan12* and *Daxx*), and generally for only 2 isoforms among the splice variants (Fig. 4A,B). These genes showed no change in overall expression upon *Otx2* knockdown (Fig. 4C). Through acute *Otx2* knockdown by viral expression of shRNA-*Otx2*, we confirmed a significant increase in the expression of only the *Mcrs-209* and *Daxx-204* isoforms, as other isoforms either did not change significantly or changed in the opposite direction (Fig. 4D). Interestingly, MCRS and DAXX interact within a protein complex with various nuclear functions including transcription regulation, chromatin remodeling, and DNA repair. Further research is needed to determine the functional consequences of these changes in distribution of transcript isoforms.

Homeoproteins have been postulated to regulate transcript splicing. The PAX6 homeoprotein can alter the population of Tenascin-C splice variants without changing total Tenascin-C expression (Von Holst et al., 2007), and the CDX2 homeoprotein interacts with splicing machinery (Balbinot et al., 2017). Regarding OTX2, its protein interactome in adult retina revealed putative RNA processing partners such as SFPQ and U2AF (Fant et al., 2015). In the present study, we found the putative RNA processing partners ACIN1, DDX41, DDX46, HNRPLL, and PRRC2A, RBM25, RBM39, SF3A1, SF3B1, SNRNP200, and SRRM2 in either ChP and/or non-cell autonomous structures (Tables 6,7). It remains to be determined whether OTX2 controls ChP splicing activity through direct interaction with splicing factors and/or by regulating their expression.

## Conclusions

Schematically, the brain can be considered as a relay organ between sensory inputs and motor actions, with the accumulation and complexification of regulatory functions through evolution resulting in extremely refined behaviors. This “getting in and getting out” relaying by the brain is not only through neural sensory and motor circuits, but also through the action of hormones, either secreted by specialized glands mainly into the bloodstream or produced by specialized neurons primarily in the hypophysis and hypothalamus. However, exchanges between the brain and the rest of the body are extremely controlled, as illustrated by the brain’s “immune privilege”, although this view has been blurred by the role of microglial cells from the immune system in early brain development (Thion et al., 2018), and by the influence of microbiota on brain development and physiology (Hsiao et al., 2013; Sherwin et al., 2019). Nonetheless, exchanges between the brain and the rest of the body are tightly regulated, and it is clear that the ChP has barrier functions for controlling what gets in and out of the brain, and homeostasis functions for controlling brain metabolites in the CSF. Our analysis reinforces important ChP endocrine functions related to OTX2. Indeed, in addition to regulating the expression and post-transcriptional modification of genes encoding signaling and hormone-transport proteins secreted into the CSF, such as *Igf2* and *Ttr*, OTX2 itself is secreted by the ChP and exerts essential non-cell autonomous activities, such as the regulation of cerebral cortex plasticity or that of adult neurogenesis. Transcriptomic analysis of different genetic *Otx2* loss-of-function models, including conditional knock-down specifically in the ChP, coupled with proteomic analysis, are first steps toward a better understanding of the molecular biology of this traveling transcription factor in and out of its main cerebral source.

## Materials and Methods

### Animal ethics statement

All animal procedures, including housing, were carried out in accordance with the recommendations of the European Economic Community (86/609/EEC), the French National Committee (87/848) and French bylaws (AGRG1240332A / AGRG1238724A / AGRG1238767A / AGRG1238729A / AGRG1238753A). For surgical procedures, animals were anesthetized with Xylazine (Rompun 2%, 5 mg/kg) and Ketamine (Imalgene 500, 80 mg/kg). For biochemical analysis, mice were either underwent transcardial perfusion or were sacrificed by cervical elongation. This research (project no. 00704.02) was approved by Ethics committee n° 59 of the French Ministry for Research and Higher Education.

### Animals and stereotaxic surgery

*Otx2^lox/lox^* mice were kindly donated by T. Lamonerie (Fossat et al., 2006) and *Otx2^+/GFP^* mice by A. Simeone (Acampora et al., 2009). Three-month old *Otx2^lox/lox^* mice were injected with Cre-Tat or vehicle as described in (Planques et al., 2019) and housed for 15 days after surgery. Adeno-associated virus (AAV) were of serotype 5 and purchased from Vector Biolabs (Malvern, USA): AAV5-CMV-EGFP; and AAV5-CMV-EGFP-U6-shRNA(mOtx2). High-titer AAV5 (~10^13^ GC/ml) were injected (2 μl per mouse) bilaterally into the LV (coordinates from bregma: x, −0.58 mm; y, ±1.28 mm; z, −2 mm) with a 10 μl Hamilton syringe (0.2 μl/min). Virus-injected mice were housed for 3 weeks after surgery. Animals were an equal mix of males and females. The *Otx2^+/GFP^* mice, littermates, and the injected *Otx2^lox/lox^* mice underwent transcardial perfusion with 20 ml phosphate buffer saline, and ChPs were dissected and processed for biochemical analysis. Virus-injected mice were sacrificed by cervical elongation for ChP extraction.

### Quantitative PCR analysis

Total RNA from LV and 4V ChPs was extracted by using the RNeasy Lipid Tissue Mini Kit (Qiagen) with DNA removal. Total RNA (10 to 20 ng) was retrotranscribed by using the QuantiTect Reverse Transcription Kit (Qiagen). Quantitative PCR (qPCR) analyses of cDNA (diluted at 1/10) were performed in triplicate with a LightCycler 480 II (Roche) using the SYBR Green I Master mix (Roche). After Tm calling verification, gene-to-*Hprt* or gene-to-*Gapdh* ratios were determined by the 2^-ΔΔCt^ method. For *Otx2* expression analysis, expression was compared to mean expression of vehicle-injected mice of the same experiment.

### RNA sequencing analysis

For analysis of conditional knockdown mice, the RNA was extracted separately from LV and 4V ChPs of Cre-Tat and vehicle-injected mice. A small sample of each ChP was tested by qPCR to ensure Cre-Tat samples had less than 50% *Otx2* expression on average compared to control mice. Duplicate samples were prepared by pooling ChP lysates from 2 x 5 Cre-Tat injected mice and from 2 x 4 vehicle-injected mice. For analysis of constitutive knockout mice, the RNA was extracted from pooled 4V ChPs of 2 x 4 *Otx2^+/GFP^* mice and 2 x 5 wildtype mice in order to obtain duplicate samples of each genotype. PolyA+ mRNA purification, mRNA sequencing, and data normalization and quantification was performed by the Genomic Paris Center (IBENS, Paris, France) using Illumina HiSeq 1500. Data are available through GEO (https://www.ncbi.nlm.nih.gov/geo) accession GSE157386.

### Isoform analysis

Raw reads were processed with fastp (Chen et al., 2018) using standard parameters, and then pseudo-aligned on mm10 gencode transcriptome using salmon (Patro et al., 2017). The quantified transcriptome was then imported in R using the IsoformSwitchAnalyzeR package (Vitting-Seerup and Sandelin, 2017; Vitting-Seerup and Sandelin, 2019) with dIFcutoff = 0.15. Isoform switch test was performed using DEXseq (Anders et al., 2012; Ritchie et al., 2015) in IsoformSwitchAnalyzeR. Gene coding potential, secondary structures, signal peptides, and protein domains were analyzed with CPAT (Wang et al., 2013), Net-Surf2 (Klausen et al., 2019), SignalP (Almagro Armenteros et al., 2019), and Pfam (Finn et al., 2014), respectively.

### Protein co-immunoprecipitation

ChP from LV and 4V from 3 months old mice were pooled and lysed with 1 ml lysis buffer (100 mM Tris pH 7.5, 1 mM EDTA, 100 mM NaCl, 1% NP40, 1 mM MgCl2, 1X protease/phosphatase inhibitor (Roche)) containing 1μl of benzonase (Roche). ChP were dissociated using 26G syringe and incubated 30 min on ice. Tubes were centrifuged at 21 000*g* for 10 min and supernatant was recovered and divided in two. Each half was incubated with 44 μg of either anti-OTX2 (ab21990, Abcam) or anti-IgG antibody (ab27478, Abcam) coupled magnetic-beads (10 mg/ml with 9.5 μg of antibody per mg of beads, Dynabeads Antibody Coupling Kit Life Tech) in lysis buffer at 4°C on rotating wheel overnight. Using magnetic separation, beads were washed 5 times in 1 mL of cold lysis buffer. Pelleted beads were eluted in 20 μl of laemmli buffer 5 min at 95 °C, and then stored at −20 °C.

For V-SVZ, RMS, and VCx, tissue from 10 mice were lysed by trituration (pipette and 26G syringe) in 10 μl lysis buffer II (20 mM Tris pH 8, 120 mM NaCl, 1% NP40, 1 mM MgCl2, 5% glycerol, 1X protease/phosphatase inhibitor) per mg of tissue supplemented with 1 μl of benzonase / 1mL of lysis buffer II. Samples were processed as described above with 25 μl of antibody coupled-beads. Pelleted beads were eluted in 30 μl of laemmli buffer.

### Mass spectrometry analysis

Proteomics analysis were performed by the Protein Mass Spectrometry Laboratory (Institut Curie, Paris, France). Eluted samples in laemmli were processed and resulting peptides were analyzed by nano-LC-MS/MS using an Ultimate 3000 system (Dionex S.A.) coupled to an Orbitrap Fusion mass spectrometer (Q-OT-qIT, Thermo Fisher Scientific). Data were acquired using Xcalibur software and the resulting Mascot files (v2.5.1) were further processed by using myProMS software (v3.9) (Poullet et al., 2007). Percolator (Spivak et al., 2009) was used for FDR calculations set to 1% peptide level. For ChP proteomics, three experiments were performed. For V-SVZ, RMS and VCx proteomics, one experiment was performed. Data are available through the ProteomeXchange Consortium via the PRIDE repository (https://www.ebi.ac.uk/pride) with identifier PXD021244.

### Ontology analysis

Genes with > 10 mean reads in at least one of the ChP samples were selected for ontology analysis. Differentially expressed gene lists were generated using threshold of *p-adj* < 0.05. Ontology term enrichment and KEGG pathways were analyzed with DAVID Bioinformatic resource v6.7 (Huang et al., 2009a; Huang et al., 2009b) and ontology terms were plotted as −log_10_ scale of the enrichment p-values. UniProt (http://www.uniprot.org) was used for obtaining functional classes (Tables 1, 3, 6, and 7). Gene list comparisons and Venn diagram data were generated with web-based tools (http://www.bioinformatics.lu/venn.php).

## Acknowledgements

This work was supported by the Région Ile-de-France DIM Cerveau et Pensée (to A.Pl.) and by the European Research Council (ERC-2013-ADG-339379 to A.Pr.). The IBENS Genomics Core facility was supported by the France Génomique national infrastructure, funded as part of the “Investissements d’Avenir” program managed by the Agence Nationale de la Recherche (ANR-10-INBS-09).

## Author Contributions

Conceptualization, A.Pl., V.O.M. and A.A.D; Methodology, A.Pl., V.O.M., C.Be., C.Bl., F.D., V.M., and A.A.D.; Software, D.B. and L.J.; Validation, V.O.M., D.B., L.J., V.M and A.A.D.; Formal Analysis, A.Pl., V.O.M, and A.A.D.; Investigation, A.Pl., V.O.M., D.B, C.Be., C.Bl., F.D. and V.M.; Resources, D.L. and A.Pr.; Data Curation, A.Pl., V.O.M., D.B., L.J., V.M. and A.A.D.; Writing – Original Draft Preparation, A.A.D.; Writing – Review & Editing, A.Pl., V.O.M., V.M., A.Pr, and A.A.D.; Visualization, A.A.D.; Supervision, D.L., A.Pr. and A.A.D.; Project Administration, A.A.D.; Funding Acquisition, A.Pr..

